# The vault particle is enclosed by a C13-symmetric cap with a positively charged exterior

**DOI:** 10.1101/2025.06.06.658390

**Authors:** Huan Li, Francesca Vallese, Oliver B. Clarke

## Abstract

Vaults are some of the largest ribonucleoprotein complexes known, and are highly conserved across eukaryotes, but both their function and key details of their architecture remain unclear. While high-resolution structures of the vault shell are available, the architecture and symmetry of the cap at either end of the vault has remained unresolved. Here we present a 2.25 Å cryo-EM structure of the vault cap, revealing an unexpected 13-fold symmetric arrangement that contrasts with the 39-fold symmetry of the vault body, with each repeating module of the cap formed by an asymmetric homotrimer of adjacent subunits, in which two C-termini remain in the vault interior, and one projects into the cytosol. The center of the cap features an unusual architecture reminiscent of the Stomatin, Flotillin and HflK/C (SPFH) superfamily, consisting of two concentric beta barrels surrounded by an interwoven two-layer stack of alpha-helices, with the innermost barrel forming a 15 Å aperture connecting the interior of the vault with the cytosol. The vault cap features a positively charged exterior and a negatively charged interior surface, with implications for binding partner recruitment and putative binding of the vault particle to microtubules and lipid rafts. These findings uncover a new facet of vault particle architecture, and have implications for engineering and design of modified vault particles for therapeutic delivery, as well as providing new opportunities for interrogating the functional roles of the vault particle in biological systems.

## Introduction

Vaults are large ribonucleoprotein complexes that are ubiquitously present in eukaryotic cells. The vault was first discovered in 1986, with an ovoid, highly regular architecture characterized by negative stain electron microscopy (*1*). The first vault structure was resolved by cryo-electron microscopy (cryo-EM) in 1999, revealing a symmetrical, barrel-like architecture (*2*). A higher resolution crystal structure conclusively resolved the overall symmetry of the vault shell, showing that it is formed by 78 copies of Major Vault Protein (MVP) arranged with D39 symmetry, but the cap regions at either end of the vault were not resolved, likely due to a symmetry mismatch with the rest of the vault shell (*3*). In addition to major vault protein (MVP) (*4*), the vault complex includes vault poly (ADP-ribose)-polymerase (vPARP) (*5*), telomerase associated protein 1 (TEP1) (*6*) and untranslated vault RNAs (vRNAs) (*7*, *8*).

Further insights into vault organization were gained through cryo-electron tomography (cryo-ET), which allowed direct visualization of vaults in intact cells. These studies showed that cargo within the vault lumen is not arranged in an ordered manner with respect to the vault walls, potentially explaining challenges in computational classification in single particle cryo-EM (*9*). Additionally, structural analysis of recombinant vaults by cryo-EM demonstrated that vault opening occurs via an expansion at the waist, which may serve as a mechanism for cargo release and uptake (*10*).

Evolutionary analysis indicates that vaults are highly conserved across multicellular eukaryotes, suggesting an ancient and fundamental cellular role (*11*), but a detailed understanding of this role remains elusive, although recent studies have suggested a role for vault in both the adaptive and innate immune response. *Mvp* knockout mice exhibited significantly attenuated immunity responses and decreased survival following homologous influenza A virus (IAV) rechallenge, in comparison to wild-type mice, suggesting that vaults can enhance adaptive immunity against viral pathogens, either directly or through interactions with innate immune pathways (*12*). Furthermore, vaults contribute to host resistance against lung infections caused by *Pseudomonas aeruginosa* (*13*).

In addition to their role in immune responses, vaults are involved in intracellular transport, particularly through interactions with the microtubule network. They have been shown to move along microtubules (*14*), and their mobility is, at least in part, dependent on microtubule integrity (*15*). Moreover, vaults associate with lipid rafts, suggesting a potential role in membrane-related cellular processes (*16*). One of the most well-characterized functions of vaults is their involvement in drug resistance in cancer. Vaults have been identified as the human lung resistance-related protein (LRP), which contributes to chemoresistance through mechanisms that are still under investigation (*17*). A recent study has shown that vaults may mitigate bone loss associated with aging and estrogen deficiency by promoting Fas-mediated apoptosis in osteoclasts. MVP prevents ubiquitination of target proteins, presumably by sequestering them from the cytosol, and MVP deficiency results in osteoporotic phenotypes *in vivo* and *in vitro* (*18*). The vault also plays significant roles in the nervous system. vRNAs directly bind MEK1, amplifying the ERK signaling pathway, which is essential for synaptic connectivity and plasticity (*19*). Additionally, vaults may be involved in nucleocytoplasmic transport, as evidence suggests they participate in the exchange of macromolecules between the nucleus and cytoplasm (*20*). Vaults have been exploited for biotechnological applications, particularly in drug delivery, owing to their large internal cavity, biocompatibility, and ability to be engineered for targeted transport. Recombinant vaults can be functionalized with targeting peptides or loaded with therapeutic molecules, enabling the delivery of drugs, nucleic acids, or proteins to specific cells or tissues with minimal immunogenicity and toxicity (*21–23*).

Despite these advances, many aspects of vault biology remain unclear. Further research is needed to elucidate their precise mechanisms of action, cargo specificity, and potential biomedical applications. Understanding these fundamental aspects will be crucial for harnessing vaults in therapeutic and biotechnological contexts. Here, using single particle cryo-EM, we show that the cap of the vault, rather than being open, as might be expected from prior structural studies, is actually a closed and tightly folded assembly, with 13-fold symmetry that contrasts with the 39-fold symmetry of the main body of the vault, and a 15 Å passage through the center of the cap that connects the vault lumen to the cytosol. These results have significant implications for our understanding of vault architecture, and for design and engineering of vaults as delivery vehicles for therapeutics.

## Results

### High-resolution structure determination of the rabbit vault particle

We purified vault from rabbit liver using two successive density gradient centrifugation steps following previously published protocols (*2*). This initial preparation yields intact vaults containing MVP, TEP1, and vPARP (Fig. S1A, S1B).

Cryo-EM analysis of the vault particles on graphene-oxide grids revealed barrel-like particles with the expected D39 apparent symmetry, yielding an initial reconstruction at 2.3 Å with D39 symmetry applied after incorporating Ewald sphere corrections to account for the significant defocus gradient across the large particle. Overall, the vault shell exhibited similar features with previously published vault structures. The vault is 700 Å long, and ∼400 Å wide (Fig. 1A). The density map was of good quality for most functional regions and we were able to fit 858 of the 890 amino acids of vault complex in the final atomic model. Residues G859 to P890, which constitute the distal C-terminus of MVP, and comprise the vault cap region, were not modelled because their densities were weak or missing, similar to previous vault structures (Fig. S1C). Despite extensive classification attempts, we were unable to identify ordered cargo within the vault lumen, consistent with prior cryo-ET results of native vaults on human umbilical vein endothelia cells (*9*).

**Fig. 1.**
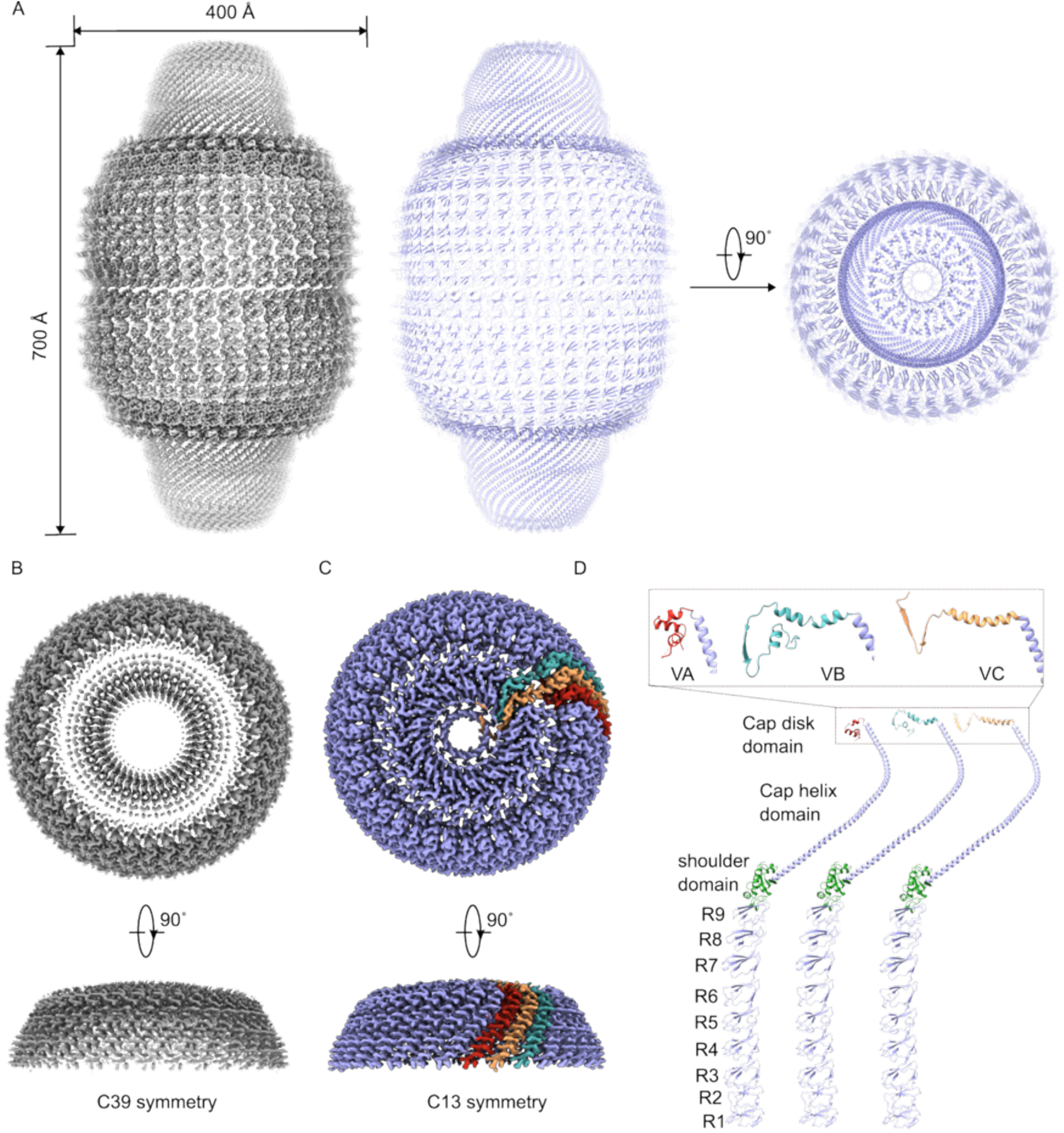
Cryo-EM structure of vault and symmetry determination of vault cap. (**A**)The overall map and structure of vault in top and side view. (**B**) The top and side view of vault cap at C39 symmetry. (**C**) The top and side view of vault cap in C13 symmetry, red, yellow, teal represent VA, VB, and VC chain, respectively. (**D**) Comparison of the conformation and secondary structure of VA, VB and VC.

To resolve the symmetry of the vault cap, D1 symmetry expansion was performed, and subparticles were extracted and reconstructed with C39 symmetry centered on the apical entrance of the vault. A block-based reconstruction approach was applied, in which the per-particle defocus values of the sub-particles was corrected based on their z-height difference with respect to the center of the vault. In C39, the walls of the vault cap had excellent density, with well resolved sidechains, but the planar cap had indistinct, disordered density that could not be confidently modeled, causing suspicion of a symmetry mismatch (Fig. 1B). Focused classification without alignments in C1, using a mask covering the planar end of the vault, revealed three classes with C13 symmetry, distinguished only by rotation about the pseudo-C39 symmetry axis. Aligning the three volumes, and adjusting the poses of the associated particles by the same transformation, facilitated local refinement with C13 symmetry, revealing the detailed architecture of the vault cap at 2.25 Å resolution (Fig. 1C).

The cryo-EM structure of the vault cap has allowed us to complete the model of the vault shell. Due to the C13 symmetry of the cap, there are three sequence-identical but distinctly folded protomers that comprise the asymmetric unit, which we have termed VA, VB, and VC. Each protomer can be divided into 12 domains from the N terminus to the C terminus: 9 structural domain repeats (R1-R9), an MVP shoulder domain, a cap helix domain, and a C-terminal cap disk domain. The cap disk domains include 3 different distinct folds in the three different protomers (Fig. 1D).

### Architecture of the vault cap

The vault shell exhibits almost perfect D39 symmetry until G803. Starting from P804, the 39 chains show 3 different folds, which weave together to form a two-layer arrangement, in which two layers of alpha-helices orthogonal to the pseudo-C39 axis buttress two concentric beta barrels, the outer of which has 26 parallel strands (contributed by VB and VC) and the inner of which has 13 parallel strands (formed solely by VC).

The three protomers have distinct folds, which come together to form the cap disk, as follows (Fig. 2A-2D). In the following description, we have numbered the secondary structural elements starting from the first helix after the cap helix domain. The VA chain is mostly α-helical (α1 to α3). VA projects partway towards the center of the cap, then VA-α2 turns outward, forming a helix-turn-helix. After a missing loop (K828 to V839), α3 stacks together with the α1 helix (Fig. 2B). 13 copies of the VA chains form an “outer ring” near the inner wall of the cap (Fig. S2A). Secondly, the VB chain contains helices α1, α2 and a single strand, β1. The curved α1 helix projects all the way to the center of cap, then the β1 strand forms the first strand in a 26-stranded barrel surrounding the apical entrance of the vault – the “outer barrel” - before emerging in the vault lumen. The luminal VB-α2 helix runs parallel to the cap disk surface (Fig. 2C). Thus, 13 copies of the VB protomer form a “middle ring” around the outline of the inner beta-barrel (Fig. S2B). Lastly, the VC protomer includes α1, β1 and β2. The VC-α1 helix extends toward the center until L825 running parallel to VB-α1, then VC-β1 projects into the vault lumen, forming the complementary strand to VB-β1, which together form the outer barrel. VC then makes a 180° turn and forms a putative beta strand, β2, which extends out back towards the cytosol and terminates at T844, forming an “inner barrel” (Fig. 2D). Overall, the cap structure looks like a sunflower when viewed from the vault lumen (Fig. S2C). VA, VB, and VC chains independently form the outer ring, the middle ring and the inner barrel, respectively, while VB and VC weave together to form the outer barrel.

**Fig. 2.**
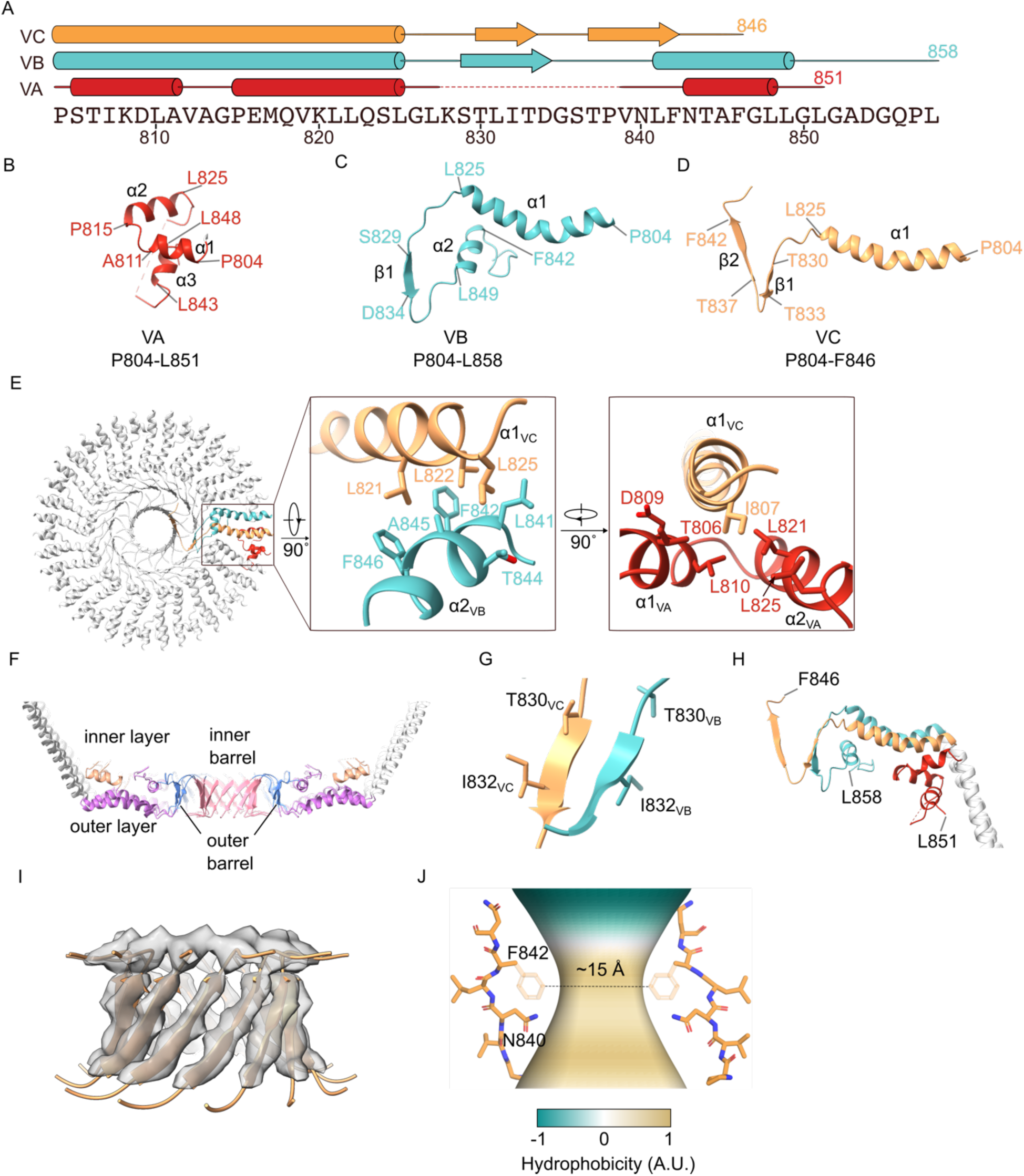
Overall architecture of vault cap. (**A**) Amino acid sequence of the vault cap. The secondary structures of VA, VB, and VC chains are indicated with cartoons. (**B-D**) The structures of VA, VB, and VC chains for vault cap. (**E**) The interactions between VA, VB, and VC chains. (**F**) The cross-section of vault cap. Outer layer represented in magenta, inner layer represented in apricot, outer barrel represented in blue, inner barrel represented in pink. (**G**) Beta strand of outer barrel, T830/I832 is adjacent in VB and VC chains. (**H**) Single VA, VB, and VC chain combination. (**I**) inner barrel structure (G835 to A845) overlay with density map; map is contoured at level 0.15. (**J**) Interior aperture of vault cap. The pore size and hydrophobicity of the barrel was calculated using software CHAP (*44*). N840 and F842 are labeled, and the sidechain of F842 (depicted with transparency) was filled using Dock Prep in ChimeraX prior to CHAP calculation.

The VC chain interacts with VA chain at the outer ring and with VB chain at the middle ring (Fig. 2E). VC chain acts like a bridge, connecting VA and VB chains. L821, L822 and L825 from VC-α1 helix are very crucial. The triple leucines insert into the VB-α2, hydrophobically interacting with L841, F842, T844, A845 and F846 from VB-α2. Regarding VA and VC chain interactions, I807 in VC-α1 helix interacts with the hydrophobic pocket in both VA-α1 and VA-α2 helices formed by T806, D809, L810, L821 and L825, respectively.

The 3 sets of chains form a two-layer, double-barrel arrangement when looking at the cross-section of vault cap. VB-α1 and VC-α1 form the outer layer. The VA outer ring and VB middle ring form the inner layer. VB and VC chains fold together to form two concentric beta-barrels in the center of the vault cap, creating a passage to the inside of the vault particle (Fig. 2F). The outer barrel consists of VB-β1 and VC-β1. Adjacent strands in the outer barrel are offset in register by one residue (Fig. 2G). The inner barrel is formed solely by VC-β2. The C-termini of both VA and VB chains project into the vault lumen, while the C-terminus of VC chain points out into the cytosol (Fig. 2H). This observation has parallels with the heterodimeric arrangement in the HflK/C complex from the SPFH family, in which the HflK subunit points inside, while HflC subunit points out to the extracellular region(*24*).

### The inner barrel forms a dynamic portal between the vault lumen and the cytosol

The inner barrel formed by the VC chain connects the vault lumen and the cytosol. In our dataset, the inner barrel is defined by 13 β-strands (Fig. 2I). These strands, while having strong density, are less well resolved than the surrounding regions, and lack most sidechain densities. It is possible that the inner barrel is flexible or dynamic, which limits the resolution in this region. Another possibility is that the inner barrel has an additional layer of pseudosymmetry which is not possible to resolve with the present dataset and currently available computational tools. It is worth noting that there is also a rod of strong density in the center of inner barrel, which could correspond either to one of the termini of the vault protomers, or potentially a small molecule or additional protein segment. Although the density for the inner barrel strands is poor, connectivity with the preceding regions of the outer barrel is clear, allowing moderate confidence in sequence assignment for VC-β2. The constriction of the inner barrel pathway is most likely formed by residues N840 and F842, with a diameter of approximately 15 Å (Fig. 2J). This constriction marks the boundary between the outward-facing hydrophilic region and the inward-facing hydrophobic region of the pathway. Consurf analysis (*25*) indicates that the inner barrel is highly conserved, especially at N840 (Fig. S2F). The 15 Å pore size appears potentially sufficient to allow diffusion of small molecules, ions, and possibly peptides. Consistent with this, strong but uninterpretable density was detected inside the inner barrel (Fig. S2G). The caveolin complex exhibits a similar structural feature, with a central barrel composed of an odd number of parallel strands (11 in the case of caveolin) (*26*).

### The vault cap forms a planar disc with a positively charged exterior and negatively charged interior

Electrostatic analysis shows that the cap has a positively charged ring formed by K828 from both VB and VC chains on the outer surface (Fig. 3A and 3C). This positively charged feature likely mediates interactions with negatively charged partners, such as nucleic acids, anionic lipids, or other proteins. K828 is broadly conserved across vertebrate species, with the exception of Xenopus and chicken, where a glutamine occupies the equivalent position, complemented by positive charges substituted in nearby regions of the cap (Fig. 3B). This substitution maintains the ability for this residue to interact with the negatively charged partners, suggesting that the ring is functionally conserved across species. The cap also has an area of negative charge on the inside (Fig. 3C). The negatively charged interior may form an electrostatic well capable of retaining or repelling small molecules or ions.

**Fig. 3.**
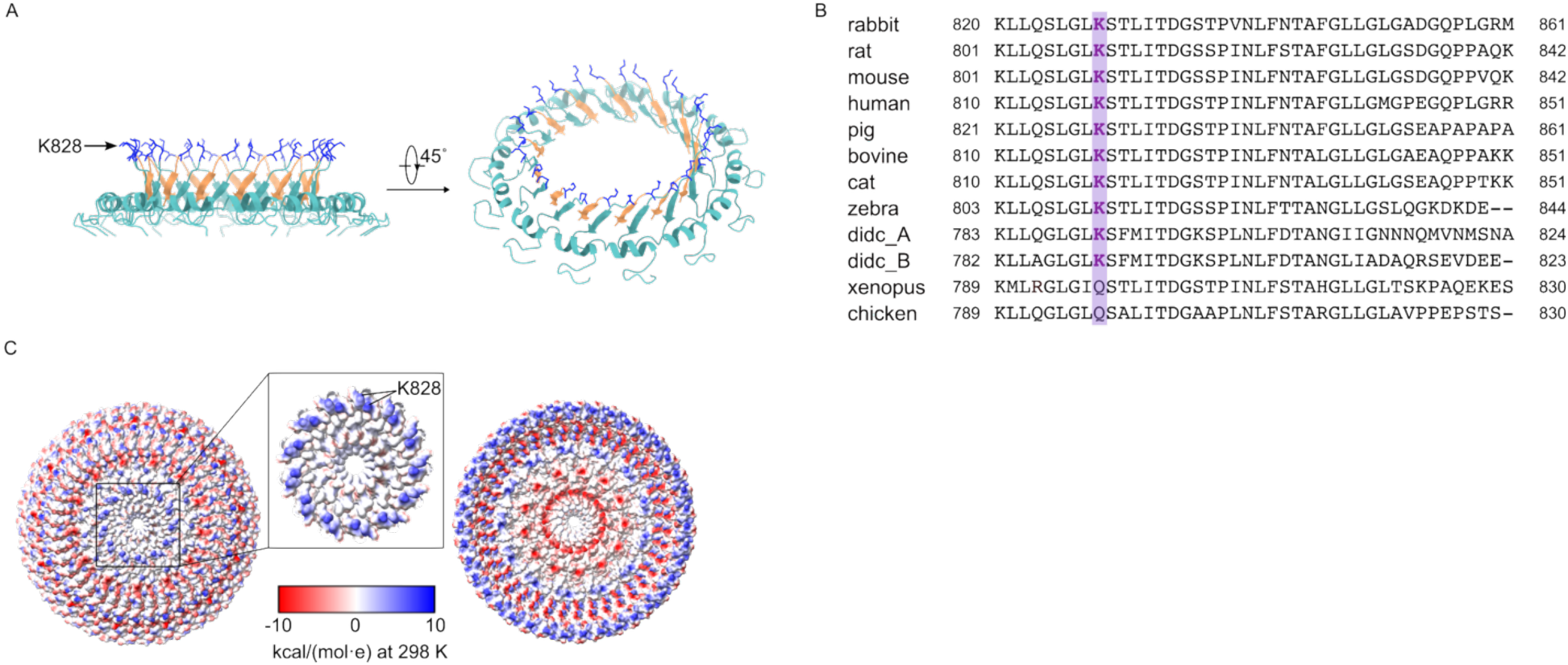
Electrostatics analysis of vault cap. (**A**) The structure of the outer barrel. K828 that forms a positive ring is shown in blue. VB chains are shown in teal and VC chains are shown in yellow. (**B**) Sequence alignment of MVP from various species, spanning residues K820 to M861 of the rabbit sequence. (**C**) Electrostatic analysis of the vault cap, shown from both exterior and interior views. Inset: zoom-in around the center of the cap from the exterior view. K828 is indicated. The color scale represents the Coulombic electrostatic potential calculated by UCSF ChimeraX, measured in kcal/(mol·e) at 298 K.

## Discussion

The vault complex is highly conserved across multicellular eukaryotes. Understanding the fundamental molecular factors that control vault dynamics is crucial, not just for unraveling the functions these particles play in various cellular processes, but also for enhancing their utility as customized molecular carriers. In this study, we report the high-resolution structure of the vault complex, revealing the architecture and symmetry of the vault cap. To our knowledge, this represents the most detailed structural analysis achieved for the vault complex to date, offering novel insights into its organization and potential functions.

Our data show that the vault cap is formed by 13 copies of an asymmetric homotrimer. While most species encode a single MVP, *Dictyostelium discoideum* (*D. discoideum*) is a notable exception, expressing two isoforms—MVP-A and MVP-B—which share 65.1% sequence similarity. This raises the possibility that *D. discoideum* vaults may form hetero-oligomers incorporating both isoforms, which would be particularly interesting given the odd number of MVP copies in a complete vault. Based on our results, the MVP shell in *D. discoideum* could adopt a 2:1 assembly, with alternating A/B/A or B/A/B subunits. Further studies will be needed to determine whether such mixed assemblies occur *in vivo*.

Previous studies have shown that vRNAs are likely bound near or within the cap and associates with TEP1, but whether the binding site is on the inside or outside has not been conclusively shown (*27*). Given the positively charged patch on the outside and the negative charge on the inside, it seems likely that vRNAs interact with the positively charged patch on the outside, if they are interacting directly with the vault cap.

### The architecture of the vault cap reveals similarities with SPFH family proteins

The cap structure of the vault particle exhibits a distinctive arrangement of two concentric β-barrels, reminiscent of architectures observed in members of the protein SPFH superfamily (Fig. 4A - 4C). A well-characterized example within this family is the HflK/C complex, whose asymmetric unit forms a heterodimer comprising one HflK and one HflC subunit. These two subunits stack to form a concentric inner barrel with 12-fold (C12) symmetry (Fig. 4B). Specifically, the outer barrel consists of 12 β-strands contributed by HflC subunits, while the inner barrel is composed of 12 strands from HflK subunits (*24*).

**Fig. 4.**
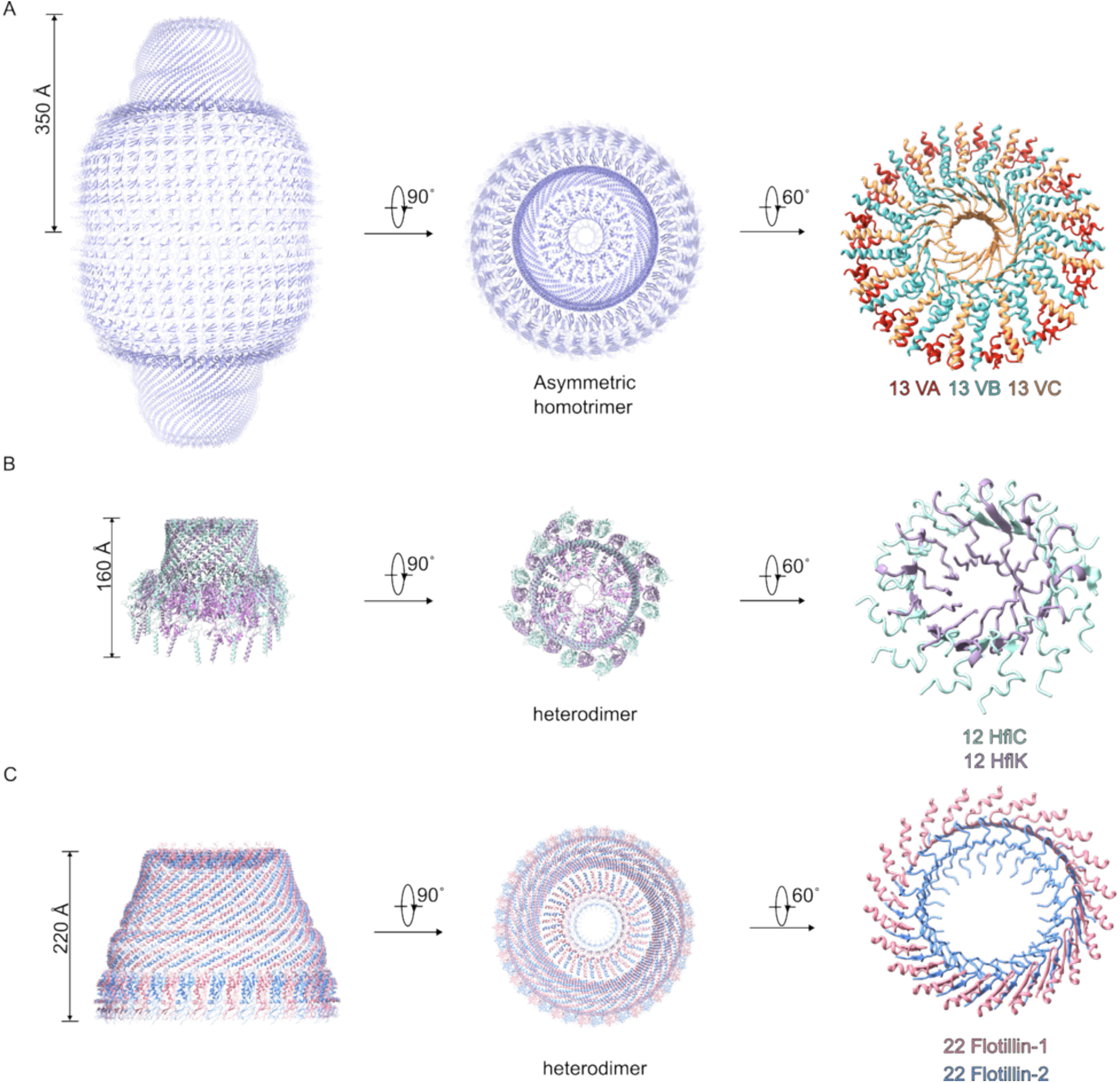
Comparison with SPFH family. (**A**) The overall structure of vault shell and zoomed in structure of cap, in three colors. (**B**) The overall structure of HflK/C complex (PDB: 7WI3) and zoomed in structure of cap, HflC and HflK in two colors. (**C**) The overall structure of Flotillin complex (PDB: 9BQ2) and zoomed in structure of cap, Flotillin1 and Flotillin2 in two colors.

A comparable structural motif is observed in the flotillin complex, in which the C-terminal regions of Flotillin 1 and Flotillin 2 assemble into a single β-barrel exhibiting C22 symmetry (Fig. 4C). Although the inner β-barrel is not explicitly modeled in the resolved structure of the flotillin complex, density map analysis suggests the possible presence of an unmodeled internal barrel (*28*, *29*). Additional members of the SPFH family, including the endoplasmic reticulum-resident Erlin complex (*30*) and the mitochondrial prohibitin complex (*31*), have recently been reported to form higher-order assemblies with similar architectural principles. In particular, the C-terminal domain of the Erlin complex forms layered structure that closely resembles the concentric organization seen in the vault cap.

While the vault cap shares these common architectural motifs, it also exhibits unique characteristics. Unlike the heteromeric assemblies typical of other SPFH family complexes, the vault shell is composed of 13 copies of an asymmetric homotrimer. Interestingly, although the three protomers are chemically identical, they adopt distinct conformations, resulting in a non-equivalent trimeric fold that contributes to the characteristic architecture of cap.

### Electrostatics of the vault cap have implications for associations with binding partners and cellular compartments

The electrostatic properties of the vault cap are essential for mediating its interactions with various cellular structures and binding partners. Notably, a prominent positively charged region is observed on the cytosolic side of the vault cap. This feature is consistent with previous observations demonstrating an end-on interaction between vault particles and microtubules, suggesting that such associations occur preferentially via the cap rather than the barrel domain of the vault (*32*). This spatial specificity in microtubule binding underscores the importance of charge distribution in directing vault positioning and trafficking within the cell.

In addition to its interaction with cytoskeletal elements, the vault cap has also been implicated in the binding of vRNAs. It has been proposed that this interaction is mediated by TEP1, which associates with both the vault and vRNAs. However, the exact binding site of vRNAs—whether internal or external to the cap—remains unresolved (*27*). The distinct electrostatic patterning of the cap offers potential insights into this question. Specifically, electrostatic surface potential mapping reveals a positively charged exterior and a predominantly negatively charged interior within the cap structure. This distribution suggests that if vRNAs interact directly with the vault cap, the interaction is more likely to occur on the outer surface, where electrostatic attraction would be maximized.

Together, these observations indicate that the electrostatic properties of the vault cap are not merely structural features but are functionally significant. They likely contribute to the selective recruitment of binding partners, such as cytoskeletal elements and nucleic acid molecules, and may influence subcellular localization and trafficking of the vault particle.

### Implications for design and engineering of modified vault particles

The structural features of the vault cap provide a framework for the rational design and engineering of modified vault particles. Its defined architecture allows for structure-guided modifications of the vault cap to enhance stability and fine-tune surface properties such as charge and hydrophobicity. Given that vault particles assemble from an asymmetric homotrimer, engineering efforts could focus on generating heterotrimeric or heterodimeric vaults with selectively stabilized conformations for each protomer. One potential approach involves truncating the segment between VA-α2 and VA-α3 to restrict flexibility, thereby favoring the VB chain conformation. Additionally, mutating G847 to alanine in VC chain may increase the α-helical propensity of this region, further stabilizing the desired structural arrangement. These modifications could provide a means to control vault assembly and improve functional specificity. Selective stabilization of conformations may enable the precise fusion of cargo within the vault interior while preserving structural integrity. Furthermore, engineered vaults could be designed to display targeting motifs or functional domains on specific protomers that project into the cytosol, while carrying cargo in the lumen. Such modifications could expand the utility of vaults in targeted drug delivery, molecular transport, or synthetic biology applications. Overall, a structure-informed approach to vault engineering has the potential to enhance their adaptability for biomedical and nanotechnological applications.

## Methods

### Purification of vaults

Vaults were purified essentially as previously described in Kong et al. (*2*). Briefly, Rabbit liver was homogenized using a polytron in buffer A containing 75 mM NaCl, 50 mM Tris-HCl, 1.5 mM MgCl_2_, 1 mM DTT, 1 mM PMSF and Pierce protease inhibitor mini tablet (Thermo scientific), with pH adjusted to 7.4. Debris was removed by centrifugation at 500 g for 15 minutes. The supernatant was passed through cheesecloth and centrifuged at 20,000g for 20 minutes. The final supernatant was centrifuged again at 100,000g for 2 h to obtain a crude microsome. The crude microsomes were then resuspended in buffer A containing 6.25% sucrose and 6.25% Ficoll-70 (Sigma Chemical Co.) and centrifuged at 40,000g for 40 minutes. The supernatant was diluted with 4 volume of buffer A and pelleted down by 100,000g for 2 h. The pellet was resuspended in buffer A with 0.5 g/ml CsCl, loaded onto a step gradient of 1.45, 1.50 and 1.70 g/ml CsCl steps, and centrifuged in a SW32 rotor for 24h at 55,000g. The load fraction was saved, diluted with 4 volume of buffer A, and centrifuged for 3h at 100,000g. The pellet was resuspended again in buffer A plus 0.5 g/ml CsCl, and loaded onto a second CsCl step gradient and fractionated as described. Final purified vaults from the load fraction were resuspended in PBS buffer containing 1 mM MgCl_2_, 0.5 mM EGTA and protease inhibitor cocktail.

### Negative stain imaging

Negative staining was performed using established methods (*33*). Briefly, 200-mesh copper grids covered with carbon-coated collodion film (Electron Microscopy Sciences, Hatfield, PA, USA) were glow-discharged for 30 s at 10 mA in a PELCO easiGlow glow discharge unit (PELCO, Fresno, CA, USA). 3 μl of purified vault sample (50 μg/ml) was adsorbed to the grids and incubated for 30 s at room temperature. Samples were then washed with two drops of water and stained with two successive drops of 0.3 % (w/v) uranyl formate (EMS, Hatfield, PA, USA) followed by blotting until dry. The sample was collected using a Tecnai Spirit T12 transmission electron microscope operated at 120 kV (Thermo Fisher Scientific, Waltham, MA, USA) at a nominal magnification of ×26,000 (2.34 Å per pixel).

### Cryo-EM data collection

3 µL purified vaults at 2 mg/mL were added to a glow discharged (PELCO EasiGlow) home-made graphene oxide (GO) grid (Quantifoil UltrAuFoil grids (R0.6/1, Au 300-mesh gold)) and blotted for 5 s at 4 °C and 100 % humidity using the Vitrobot Mark IV system (TFS), before plunging immediately into liquid ethane for vitrification. For home-made GO grids, 2.4% (w/v) GO solution (Sigma) dissolved in 20% ethanol solution was vortexed and centrifuged at 3000 rpm for 5 minutes. 1 µL of the supernatant was added on the top of each grid and allowed to air dry. Images were acquired on a Titan Krios electron microscope (TFS) equipped with a K3 direct electron detector (Gatan) and a Gatan Image Filter (GIF) operating at 64000x magnification with 0.68 Å per pixel in super-resolution mode (1.34 Å physical pixel size) using Leginon automated data collection software (*34*, *35*). Data collection was performed using a dose of 54 e^−^/Å^2^ across 50 frames (100 ms per frame) at a dose rate of 14 e^−^ /pixel^−1^ s^−1^, using a set defocus range of −0.5 to −1.5 μm. A 100-μm objective aperture was used. Data was collected using image shift with hardware coma correction. A total of 11,213 micrographs were collected.

### Cryo-EM data processing

The brief cryo-EM data processing workflow is summarized in Fig. S3. Euler angle distributions, conical Fourier shell correlation (cFSC) plots and validation statistics are presented in Fig. S4 and Table 1. Maps, masks and raw videos have been deposited at EMDB (Table 1). Subsequent steps were performed in CryoSPARC (*36*) unless otherwise indicated.

**Table 1:** Cryo-EM data collection, refinement and validation of vault.

|  | overall vault<br>(EMDB-TBA)<br>(PDB-TBA) | vault cap<br>(EMDB-TBA)<br>(PDB-TBA) |
| --- | --- | --- |
| **Data collection and processing** |  |  |
| Magnification | 64000 |  |
| Voltage (kV) | 300 |  |
| Electron exposure (e-/Å<sup>2</sup>) | 54.65 |  |
| Defocus range (µm) | -0.5 to -1.5 |  |
| Pixel size (Å) | 0.68 |  |
| Initial micrographs (no.) | 11213 |  |
| Final micrographs (no.) | 7319 |  |
| Symmetry imposed | D39 | C13 |
| Initial particle images (no.) | 138,437 | 159,668 |
| Final particle images (no.) | 79,584 | 129,638 |
| Map resolution (Å) | 2.41 | 2.25 |
| FSC threshold | 0.143 | 0.143 |
| **Refinement** |  |  |
| Initial model used (PDB code) | 4HL8 and 7PKR |  |
| Map sharpening *B* factor (Å<sup>2</sup>) | 73.6 | 68.1 |
| Model composition |  |  |
| Non-hydrogen atoms | 18904 |  |
| Protein residues | 2405 |  |
| *B* factors (Å<sup>2</sup>) |  |  |
| Protein | 121.18 |  |
| R.m.s. deviations |  |  |
| Bond lengths (Å) | 0.005 |  |
| Bond angles (°) | 1.107 |  |
| Validation |  |  |
| MolProbity score | 1.96 |  |
| Clashscore | 3.66 |  |
| Poor rotamers (%) | 4.21 |  |
| Ramachandran plot |  |  |
| Favored (%) | 95.25 |  |
| Allowed (%) | 4.75 |  |
| Disallowed (%) | 0.00 |  |

Movie frames were aligned with Patch Motion job type in CryoSPARC. After CTF estimation with Patch CTF, low quality micrographs with high relative ice thickness, high motion pixel distances, and low CTF fit resolution were removed from each dataset. Vault particles were picked using template-based particle picker in CryoSPARC using templates derived from initial 2D classes. Low-resolution ab-initio models were generated with the initial sets of particles that went through a round of 2D classification. Iterative rounds of 2D classification and heterogeneous refinement further removed particles that do not contribute to high-resolution reconstructions. A second round of particle picking was conducted with Topaz, using particles manually curated from the initial selection to train the neural network (*37*). After cleaning up the particle sets for each dataset, particles were extracted with a box size of 700-pixel size (px), global consensus reconstructions of vault were carried out with Non-Uniform (NU) refinement in CryoSPARC (*38*) with D39 symmetry enforced. The particle stacks from NU refinement were subjected to reference-based motion correction (RBMC) in CryoSPARC. After RBMC, particles were grouped into 25 to 50 image shift groups to enable grouped refinement of beam tilt and trefoil aberrations. The pixel size was calibrated by fitting the vault crystal structure to the map at a range of nominal pixel sizes using fitmap in UCSF Chimera, and identifying the pixel size resulting in the maximum cross-correlation.

In the global homogeneous and non-uniform (NU) refinement maps, the cap region of the vault appeared poorly resolved, with smeared or uninterpretable density. To improve the local resolution of this region and resolve the local symmetry, local processing was performed as follows. First, local soft masks were generated using UCSF ChimeraX (37) and CryoSPARC. Then, particles were symmetry-expanded using D1 symmetry. Subparticles were then extracted in a 256 px box centered on each cap region, and the per-subparticle defocus was adjusted based on the calculated z-height difference with the center of the vault using a CryoSPARC Tools script. These subparticles were then subjected to local refinement with C39 symmetry using the cap-region mask. Focused classification on the cap region without alignment in C1 revealed 3 C13-symmetric classes distinguished by rotation around the z-axis. The 3 classes were aligned using Align 3D maps in CryoSPARC, updating particle alignments accordingly, and subjected to local refinement using the cap-region mask, with C13 symmetry enforced. Additional 3D classification was performed and one final round of local refinement with C13 symmetry was applied to the best class.

### Atomic model building and refinement

Published vault structures (Protein Data Bank (PDB) ID 4HL8 (*39*) and PDB 7PKR (*10*) were used as initial models for building. The atomic structures of vault were fitted in cryo-EM maps, and the cap region was then manually extended and completed in Coot (*40*). The structures and EM maps were visualized using UCSF Chimera (*41*) or ChimeraX. The model for the trimeric asymmetric unit was expanded into a 13x3-mer using SERVALCAT (*42*), followed by the refinement of the model using Phenix Real Space Refinement (Fig. S5). In order to aid model building of the overall vault, composite maps were generated by aligning vault cap and vault shell maps to the global reconstruction, and then combining them by taking the maximum value at each voxel using UCSF Chimera. Finally, models were fit to the composite map, merged, then flexibly fit using Phenix Real Space Refinement. Side chains of F842 and N843 were removed because of poor density. Dock prep (*43*) was performed for electrostatic and CHAP analysis. Figures were prepared using UCSF ChimeraX. Refinement and validation statistics are provided in Table 1.

## Acknowledgements

We thank members of the Columbia University Cryo-Electron Microscopy Center for assistance in cryo-EM grid screening and data collection. We would also like to thank Zhengshan Hu for manuscript feedback and support.

## Supplementary Figures

**Fig. S1.**
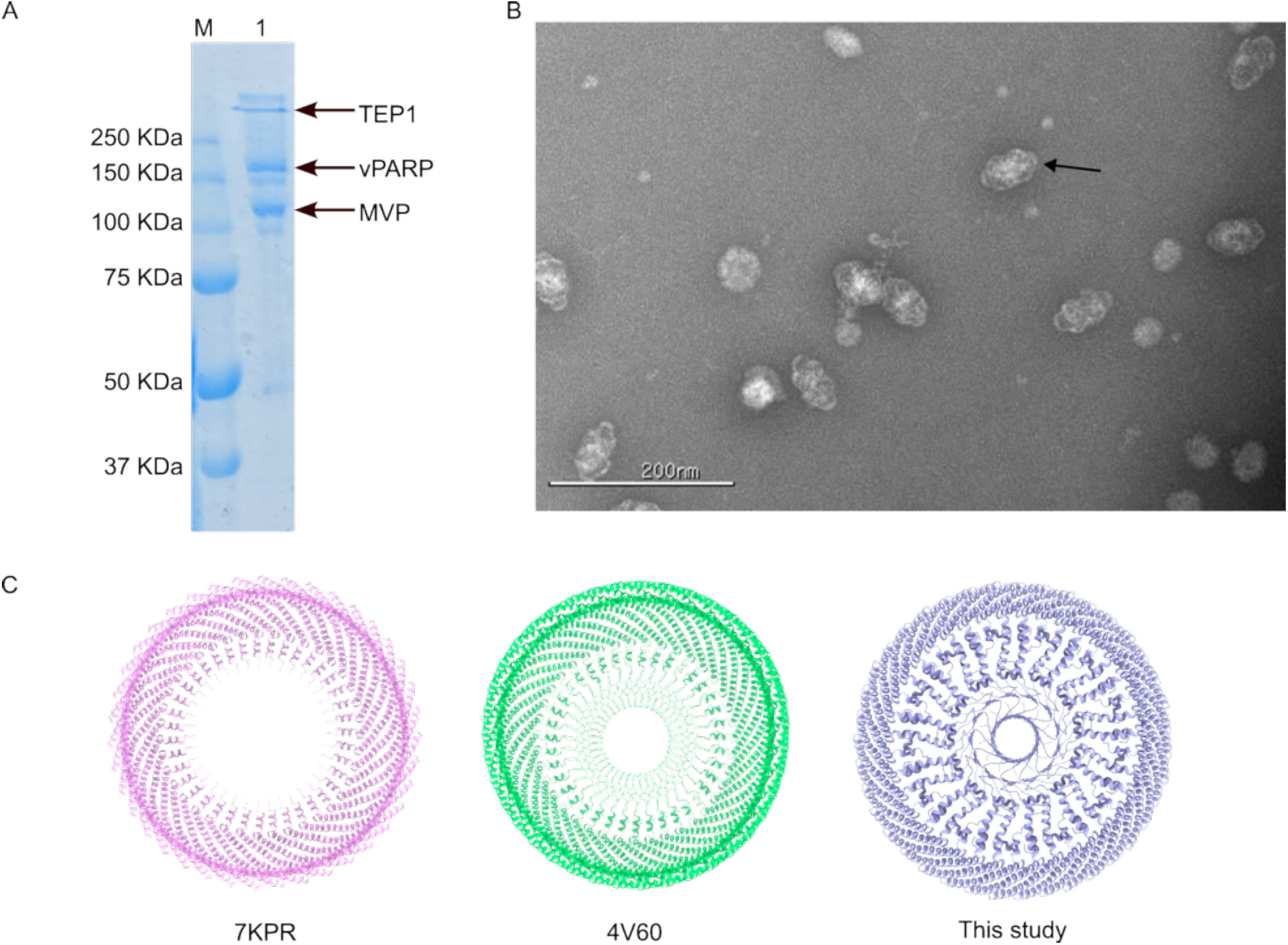
Purification and characterization of vault. (**A**) SDS-PAGE of purified vault. M: protein marker; 1: purified vault. (**B**) Negative staining of purified vault. The representative vault was indicated by arrow. (**C**) Comparison of vault cap structures.

**Fig. S2.**
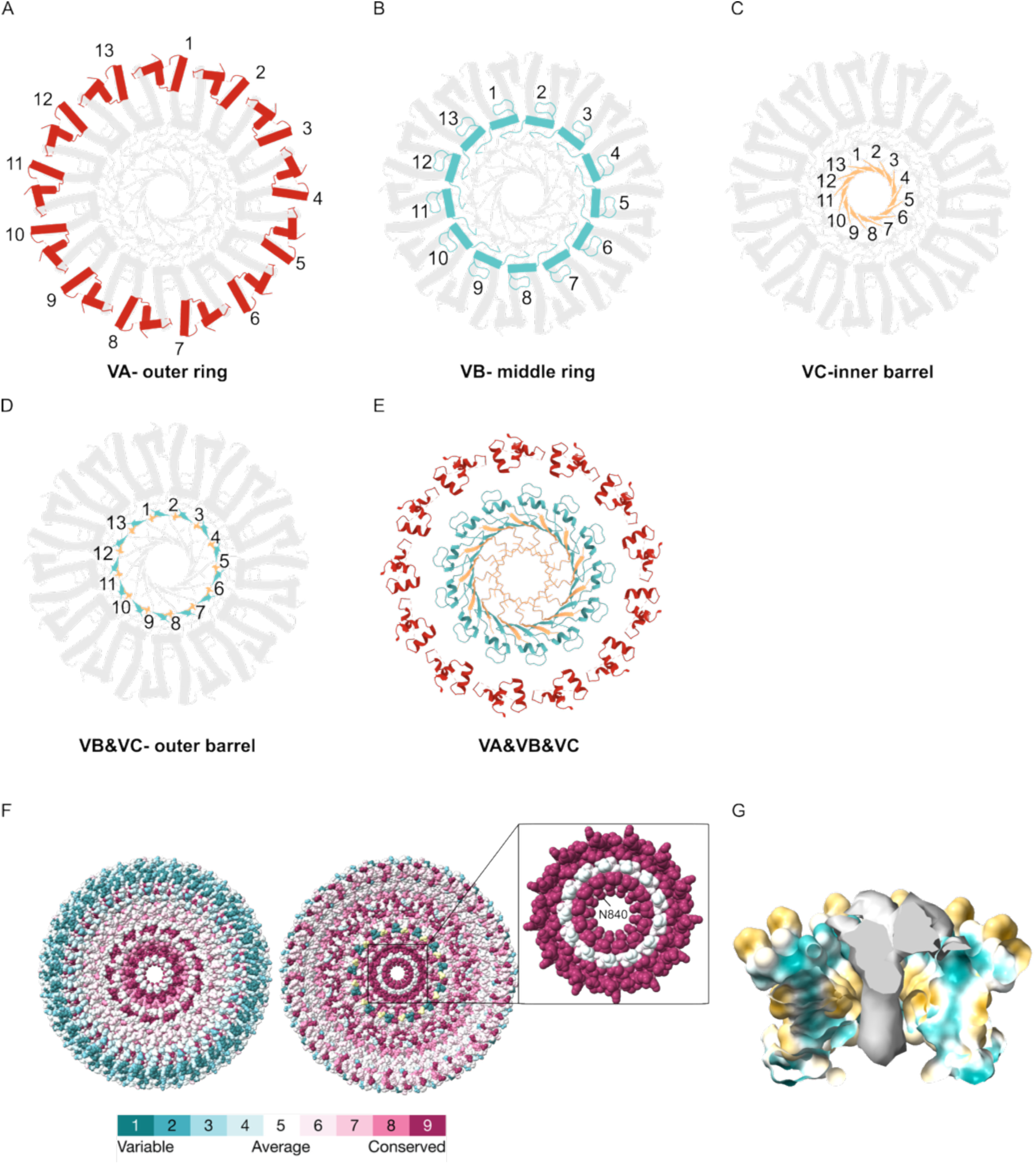
The VA, VB, and VC rings of cap. (**A**) Outer rings formed by VA chains in red. (**B**) Middle rings formed by VB chains in teal. (**C**) Inner barrel formed by VC chains in yellow. (**D**) Outer barrel was formed by partial VB and VC chains in teal and yellow, respectively. (**E**) The top view of outer ring, middle ring and inner barrel combined. (**F**) The Consurf analysis of vault cap viewed from the lumen (left) and cytosol (right). Inset: zoom-in around the center of the cap from the inside of view. N840 is labeled. (**G**) Identified cryo-EM density in the center of inner barrel. The map is contoured at level 0.168.

**Fig. S3.**
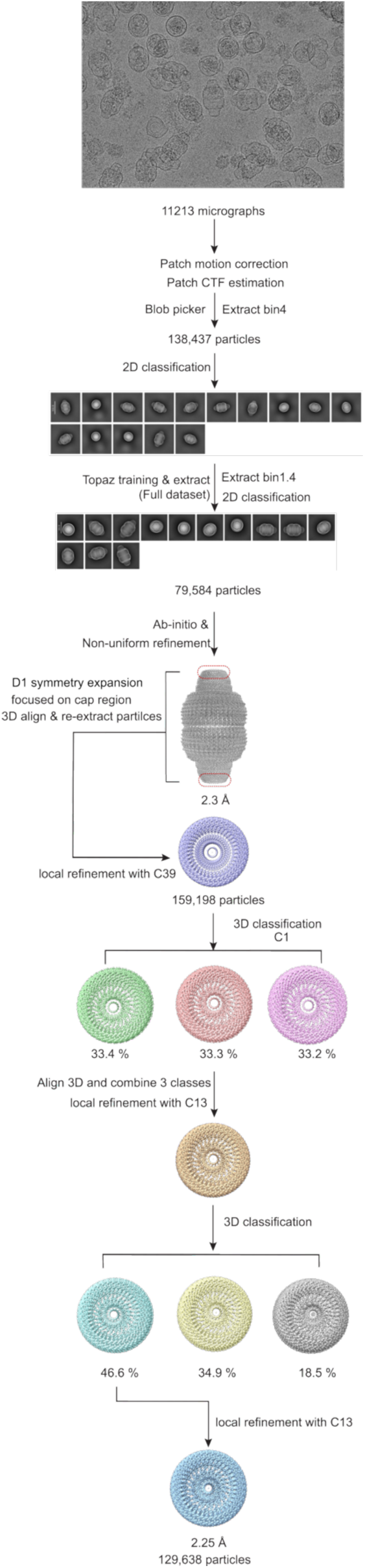
Flow chart of cryo-EM processing steps. Flowchart outlining cryo-EM image acquisition and processing performed to obtain the structure of overall vault and vault cap.

**Fig. S4.**
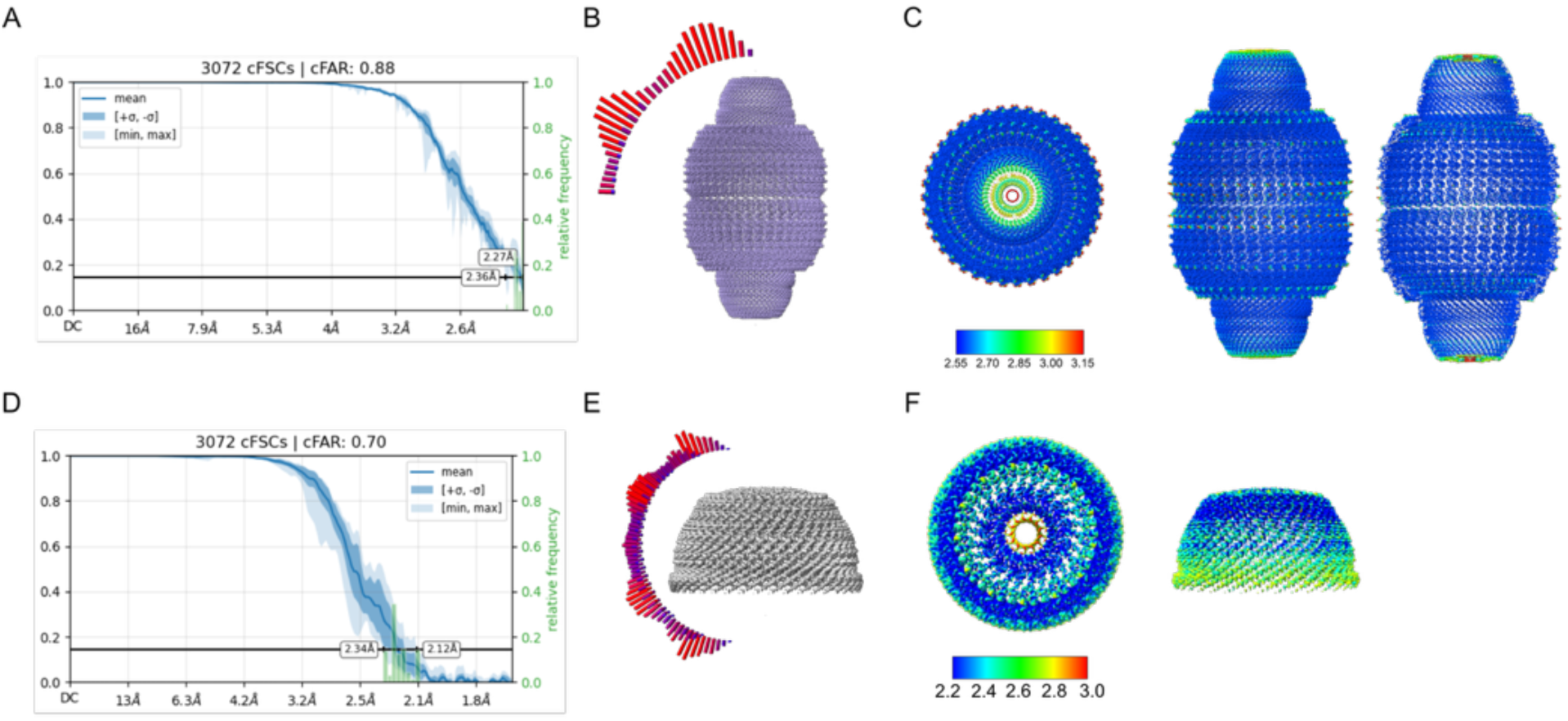
Quality of vault maps and models. (A) Conical Fourier shell correlation (cFSC) vs. resolution curves for the cryo-EM maps of vault. (B) Euler angle distribution of particles used in the overall vault map. (**C**) Top (left), side (middle), and clipped (right) viewc map of vault. (**D**) Conical Fourier shell correlation (cFSC) vs. resolution curves for the cryo-EM maps of vault cap. (**E**) Euler angle distribution of particles used in the cap map. (**F**) Top (left), side (right) views of the cryo-EM map of vault cap.

**Fig. S5.**
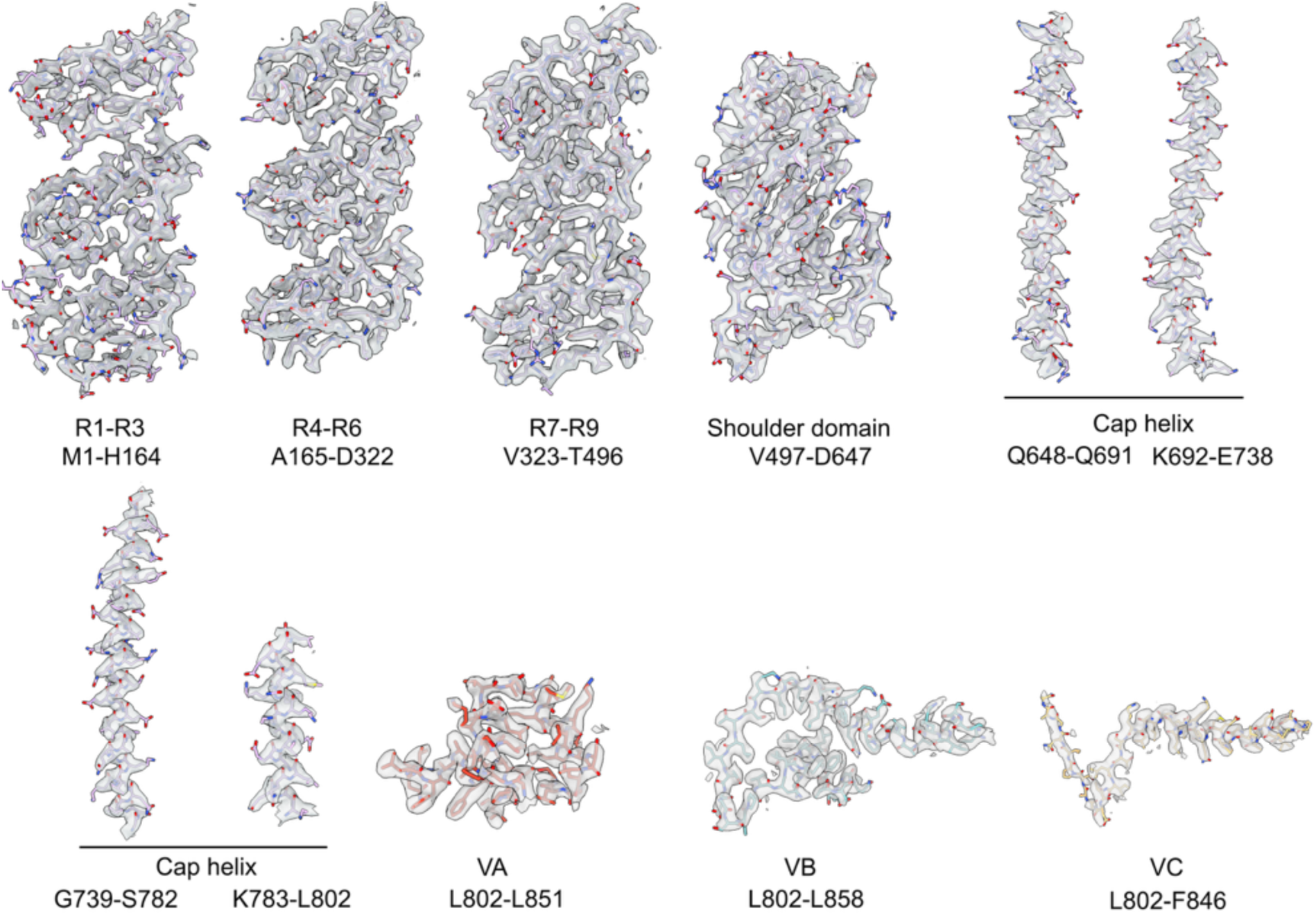
Examples of density fits of the MVP shell atomic model. Cryo-EM densities (transparent gray surface) are shown with corresponding segments of the atomic model. The composite map is contoured at level 0.18.

